# Context-dependent structurally informed effective connectivity under psilocybin

**DOI:** 10.1101/2025.08.18.671000

**Authors:** Matthew D. Greaves, Tamrin Barta, Leonardo Novelli, Devon Stoliker, Adeel Razi

**Author notes:** joint first author (Matthew D. Greaves, Tamrin Barta).

## Abstract

The extent to which anatomical connectivity constrains pharmacologically altered brain dynamics remains poorly understood. Here, we combined psilocybin administration with a structurally informed effective-connectivity model to examine how structural connectivity shapes directed inter-regional influences across experiential contexts. Using dynamic causal modeling embedded in a hierarchical empirical Bayes framework, we analyzed fMRI data acquired from a hippocampo–thalamo–cortical network during rest, guided meditation, music listening and movie viewing. Across contexts, psilocybin reorganized directed interactions while preserving structure-based scaling. Effects converged on efferents (outgoing influences) from the left hippocampus—a hub interfacing mnemonic and associative systems with the default-mode network and thalamus. Notably, the left-hippocampus-to-thalamus pathway showed a sign-reversed association with mystical-experience scores (downregulation during guided meditation and upregulation during music listening). In model-based leave-one-out cross-validation, left-hippocampal efferents predicted individual differences in mystical-experience intensity. A minimal model-free benchmark (hippocampal signal variability) also showed modest associations with mystical experience. Together, these findings link context-specific, structurally informed effective connectivity to individual differences in the acute psychedelic experience, providing a mechanistic bridge between anatomy, neurodynamics, and phenomenology.

## Main

Classic psychedelic compounds such as psilocybin, profoundly alter perception, cognition, and emotion, producing subjective experiences that can include vivid sensory phenomena, an altered sense of self, and sustained positive affect^1,2^. Recently, neuroimaging has revealed that these psychedelic effects are accompanied by widespread changes in neural dynamics, including altered large-scale network organization^3^, increased signal variance^4^, and shifts in spectral power^5^. Although both domains have garnered substantial research attention, reconciling the psychological and neurophysiological effects of psychedelics remains an open challenge.

Theoretical accounts of psychedelic action seek to bridge these psychological and neurophysiological observations in terms of a relaxing of constraints in the brain^6,7^. In this view, pharmacological action at serotonin 5-HT_2A_ receptors destabilizes entrenched patterns of neuronal communication and broadens the repertoire of accessible brain states. From a psychological perspective, this broadening has been framed as a loosening of rigid beliefs or cognitive biases—often cast as ‘priors’, in line with Bayesian terminology^6^. From a neurophysiological perspective, this broadening is expected to manifest as a partial decoupling of functional dynamics from the brain’s anatomical scaffold (in other words, weaker alignment with the brain’s white-matter topology and weights^8^).

Empirical work has begun to examine these ideas, with at least one study reporting reduced structure– function coupling under psychedelics (as inferred using a popular eigendecomposition procedure^9^). However, most research examining brain dynamics under psilocybin comes from small, heterogeneous samples^10^ and often relies on analyses of functional connectivity (statistical dependencies between observed neural signals), which, while informative, cannot resolve directed (source-to-target) communication in the brain^11^. By contrast, effective connectivity represents directed, signed influences between neuronal populations under a generative (state-space) model, enabling mechanistic, circuit-level interpretation beyond what correlation-based measures afford. Furthermore, with a few important exceptions^12–14^, neurodynamic changes under psilocybin are typically assessed in a single resting-state condition, leaving it unclear whether observed patterns generalize across diverse experiential contexts.

Here, we leverage a large, single-site functional MRI (fMRI) dataset comprising multiple experiential contexts—resting state, guided meditation, music listening, and movie viewing—acquired from the same individuals in an open-label trial under both baseline (drug-free) and psilocybin conditions (Fig. 1a–d)^15,16^. Using dynamic causal modeling and a hierarchical (parametric) empirical Bayes framework that explicitly links structural (or anatomical) connectivity to the variance (or noisiness) of effective connectivity (via priors, Fig. 1e–j), we examine how this variability is constrained by anatomy and explore psilocybin-modulated alterations to this relationship^17^. This framework also enables out-of-sample tests of whether context-specific, structurally informed changes in effective connectivity track individual differences in the intensity of the psychedelic experience.

**Fig. 1.**
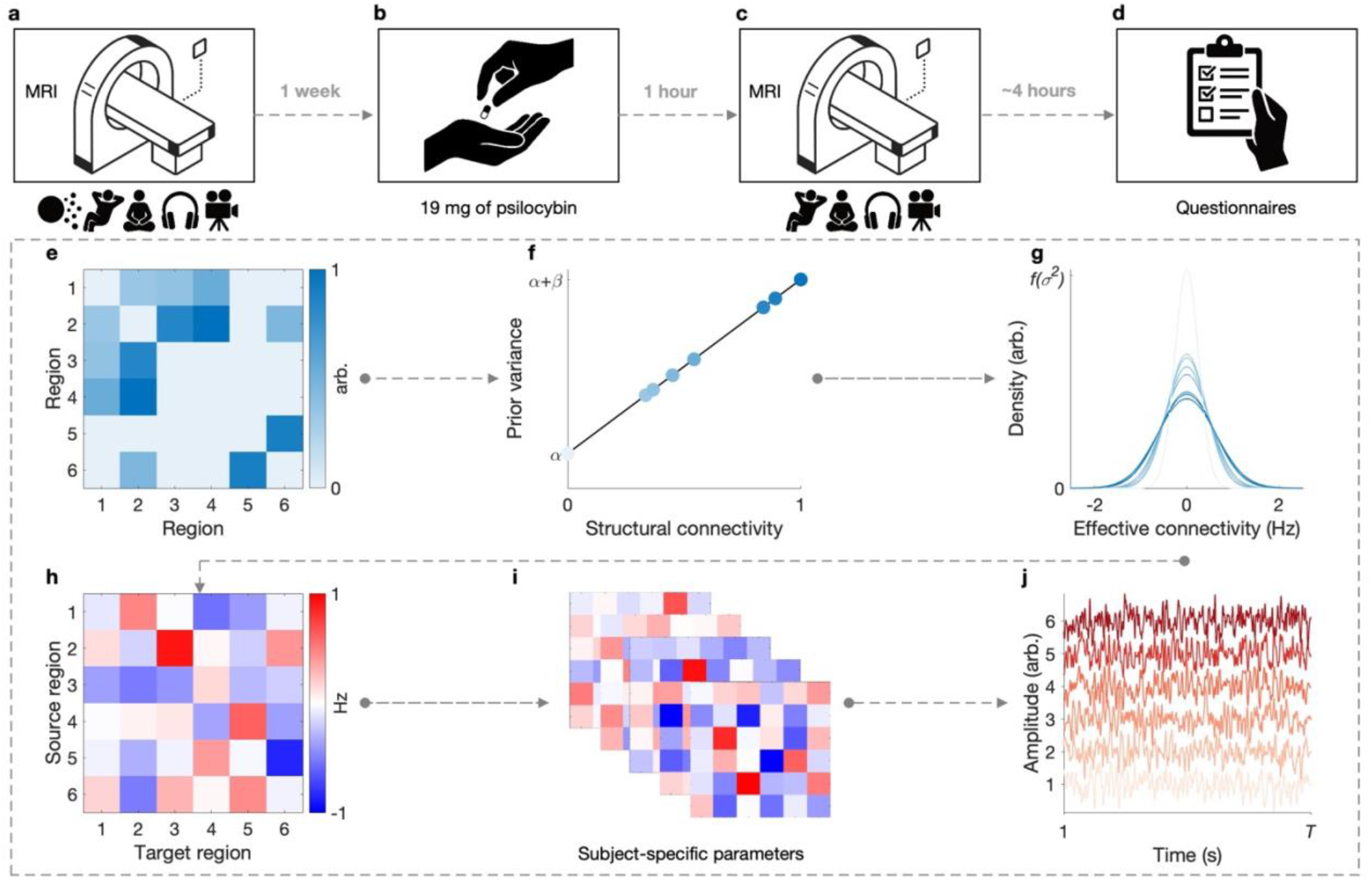
Study design and hierarchical empirical Bayes modeling framework. (**a–d**) Study protocol (excerpt). (**a**) Baseline MRI session comprising diffusion MRI, resting-state functional MRI (fMRI), and fMRI during guided meditation, music listening, and movie viewing (icons beneath scanner depict each modality/task in order: diffusion MRI, resting-state, meditation, music listening, and silent movie viewing). (**b**) Oral administration of 19 mg psilocybin (occurring approximately 1 week after baseline). (**c**) Second MRI session (1 hour after drug administration), with the same sequence of fMRI scans. (**d**) Questionnaire-based assessments completed approximately 4 hours post-administration. (**e–j**) In the hierarchical empirical Bayes model, prior variances on second-, group-level effective connectivity parameters are parameterized as a linear function of (normalized) group-level structural connectivity weights (**e**), with intercept *α* and slope *β*. When *β* > 0, this relationship implies broader priors for connections with greater structural connectivity, as shown by the color-coded scatters (**f**). Structurally informed prior variances then constrain zero-mean Gaussian distributions over connection strengths (**g**). From a generative modeling perspective, group-level effective connectivity (**h**) is sampled from structurally informed priors, and subject-specific effective connectivity (**i**) represent noisy deviations from this group connectivity profile. Simulated blood-oxygen-dependent (BOLD) time series for a single subject in the model are shown in (**j**). In the context of this study, the hierarchical model was inverted using a cross-spectral density formulation of dynamic causal modeling^18^.

We show that psilocybin reorganizes directed interactions within a hippocampo–thalamo–cortical network while preserving structure-based scaling, and that these changes are context sensitive. Across conditions, effects concentrated on left-hippocampal efferents (outgoing influences). Using leave-one-out cross-validation, single-connection left-hippocampal models predicted individual differences in mystical-experience scores. Hippocampal signal variability—a benchmark for minimal, non-mechanistic information—likewise showed modest associations. Together, these results link context-specific, structurally informed effective connectivity to individual differences in the acute psychedelic experience, providing a mechanistic bridge between anatomy, directed network dynamics, and phenomenology.

## Results

In this study, we utilized both fMRI and structural connectivity data (derived from tractography applied to diffusion-weighted MRI) from 61 healthy adults (28 female, mean age = 37.30, s.d. = 10.90) collected as part of the PsiConnect trial^15,16^. All subjects were psychedelic-naive (had no prior experience with classic psychedelic substances), except one who reported previous use (on one occasion) with no psychedelic effect. Each subject underwent scanning before and after receiving a 19 mg oral dose of psilocybin, with fMRI data acquired during resting state and three naturalistic conditions: guided meditation, music listening, and movie viewing. Apart from movie viewing, subjects kept eyes closed in all other contexts. Prior to viewing the movie (a video of moving clouds without audio), subjects were asked to make a mental note of any structured imagery (faces or repeating patterns, for example), to facilitate the reporting of these impressions (on a psychometric measure) post scan. Following the psilocybin session, subjects completed the revised mystical experience questionnaire (MEQ)^19^, providing a quantitative indicator of the intensity of their subjective psychedelic experience.

### Baseline–psilocybin changes in structurally informed effective connectivity

In our analysis, we considered a single brain network comprising regions previously implicated in psychedelic action: the medial prefrontal cortex (MPFC), posterior cingulate cortex (PCC), bilateral anterior insular (AI), bilateral hippocampus, and thalamus^3,10,20,21^. Fig. 2a shows a graph-based representation of the group average structural connectivity network projected on a cortical mesh (where line weight is proportional to connection strength). Note that, in line with previous investigations^22,23^, the left–right asymmetry in structural connectivity between the thalamus and hippocampus was present for most subjects (after correcting for spatial and geometric confounds; SI Methods).

**Fig. 2.**
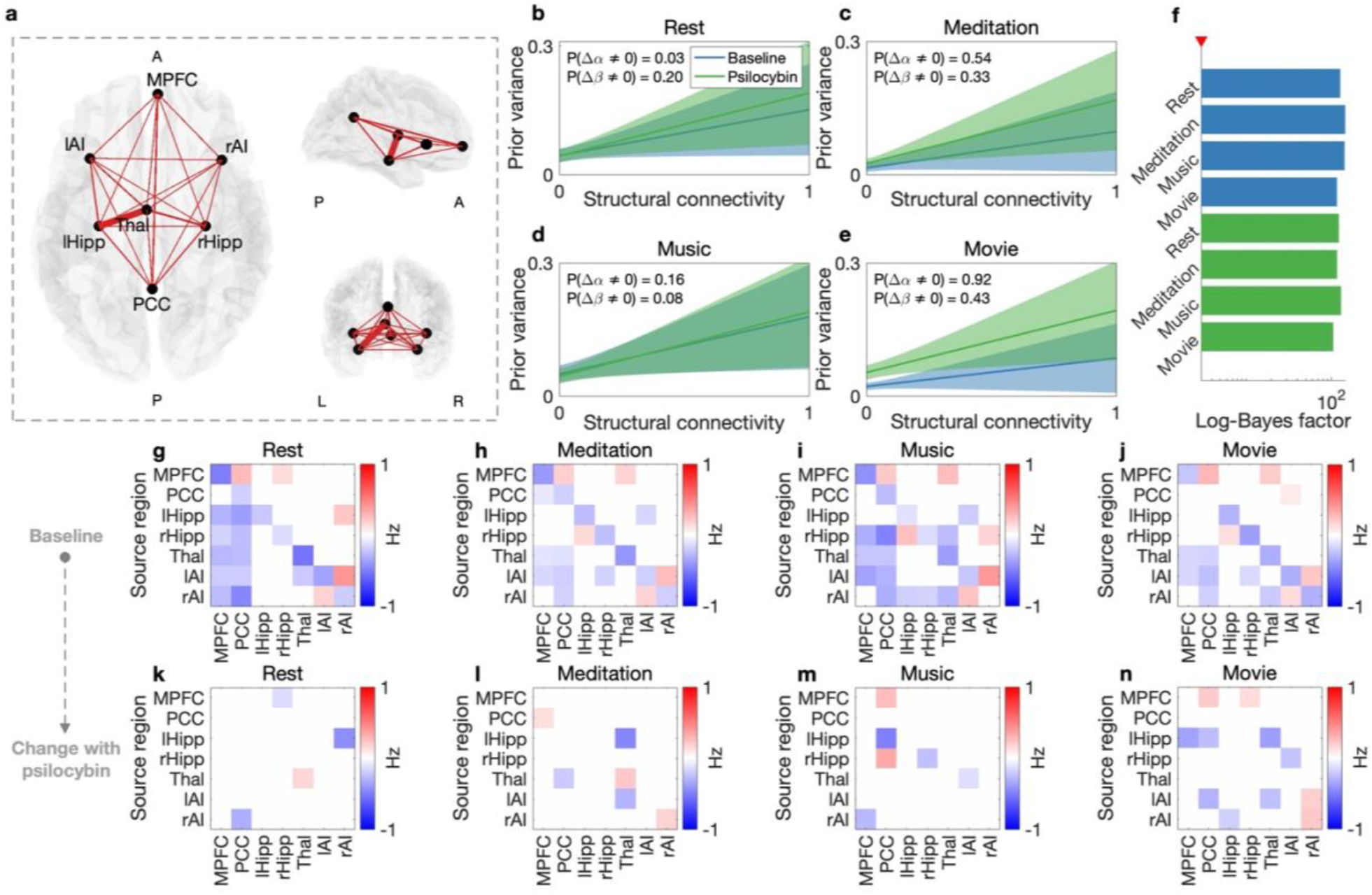
Structural connectivity constrains psilocybin-modulated effective connectivity. **(a)**Graph depiction of the considered brain networks (normalized) structural connectivity projected on a cortical mesh (axial, sagittal, and coronal views). Line weight is proportional to connection strength. Network comprised the medial prefrontal cortex (MPFC), posterior cingulate cortex (PCC), bilateral anterior insula (lAI, rAI), bilateral hippocampus (lHipp, rHipp), and thalamus (Thal). **(b.e)** Evidence-weighted prior-variance transformations show how structural connectivity scales prior variance of effective connectivity across contexts. Shading denotes standard error envelope. Results were broadly consistent across contexts when comparing prior variance at baseline (blue) and under psilocybin (green), with hyperparameters.intercept ()) and slope (β)_stable except for an intercept shift in the movie-viewing condition. Two-tailed posterior probability of baseline.psilocybin parameter differences reported in top left of each panel. **(f)** Log-Bayes factors showing increase in group-level evidence seen when each hierarchical empirical Bayes model was inverted under its evidence-weighted prior-variance transformation (green: psilocybin; blue: baseline). Note red triangle indicates log-Bayes factor of 3 (x-axis is log scaled). Subject-level log-Bayes factors not shown (SI Results, Fig. S1). **(g.j)** Group-level effective connectivity at baseline (maximum a posteriori estimates are thresholded at 90% posterior probability). Red indicates excitation and blue indicates inhibition. Note that self-connections (diagonal entries) are scaled relative to a fixed inhibitory prior (.0.5 Hz) via a log-normal transformation; values reflect relative changes, not absolute Hz. **(k.n)** Additive changes to effective connectivity under psilocybin (summed matrices depicting the full psilocybin profiles, SI Results, Fig. S2).

Per our prior methodological work, to explore the role of structural connectivity in shaping effective connectivity under psilocybin, we first inverted a hierarchical empirical Bayes model for each session (baseline and psilocybin), and context, under uninformative priors (Methods). Then, focusing on the second-, group-level model, we utilized a grid search and Bayesian model reduction (BMR) to evaluate *k* = 1, …, *K* different parametrizations of a linear structural-connectivity-to-prior-variance transformation (henceforth referred to as the prior-variance transformation) and score the resultant (structurally informed) reduced models against the (uninformed) full model in terms of the log-Bayes factor (Methods)^24^. We then applied Bayesian model averaging (BMA)^25^, weighting the hyperparameters 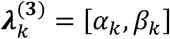, governing the *k*-th transformation by the resultant (normalized) model evidence, thereby yielding an evidence-weighted prior-variance transformation (defined by intercept *α*, and slope *β*).

Fig. 2b–e illustrates that, despite the evaluation of hyperparameter regimes resulting in flat prior-variance transformations (*β* = 0, disregarding structural connectivity), the resulting evidence-weighted prior-variance transformation for each session and context was a positive, monotonic function. This indicates that, on average, reduced models in which structural connectivity scaled the variance of effective connectivity were better supported by the data. The evidence-weighted prior-variance transformation led to a substantial increase in evidence—quantified via the log-Bayes factor—at both the group (Fig. 2f) and subject level (SI Results, Fig. S1), well beyond the conventional benchmark of 3 (strong evidence^26^). Across sessions and contexts, slopes (*β*) were stable, whereas the intercept (*α*) increased under psilocybin only for movie viewing, indicating a context-dependent rise in overall variance without a change in structure-based scaling (posterior probabilities, Fig. 2b–e). Thus, although structural connectivity did not appear to exhibit greater or lesser influence on the variance of effective connectivity under psilocybin, the baseline variance of effective connectivity did nevertheless increase under the drug (in line with prior work showing eyes-closed–eyes-open differences in the variance of functional connectivity under psilocybin^16^).

Leveraging the structural connectivity-based priors (henceforth structure-based priors) furnished by the session-specific prior-variance transformations (Fig. 2a–e), we inverted hierarchical empirical Bayes models in which baseline–psilocybin effective connectivity changes were quantified (Methods). Fig. 2g–h shows the baseline effective connectivity after thresholding maximum *a posteriori* (MAP) estimates according to the (two-tailed) 90% posterior probability of being non-zero. Across contexts, bilateral AI regions exhibited reciprocal excitation (positive-valued effective connectivity), while both the thalamus and bilateral AI inhibited PCC and MPFC default-mode hubs. In line with theoretical expectations^27^, inhibition from lower- and intermediate-order regions to these hubs was altered in the guided meditation and movie-viewing contexts (where subjects received explicit instructions), relative to task-free contexts.

Fig. 2k–n show (thresholded) additive changes in structurally informed effective connectivity under psilocybin (summed matrices depicting the full psilocybin profiles, SI Results, Fig. S2). Across contexts, the consistent features were alterations to efferents from left hippocampus and right AI (including changes to its self-connectivity), in addition to altered afferents (incoming influences) to the PCC. Beyond this, the pattern of changes was context dependent. Contexts with an auditory component (guided meditation and music listening) showed altered thalamic influences consistent with bottom-up downregulation of efferents from the thalamus to PCC and to left AI, respectively, whereas task-based contexts (guided meditation and movie viewing) were characterized by increased top-down inhibition of the thalamus from left AI. The eyes-open movie-viewing context showed the broadest set of alterations, with prominent changes in left-hippocampal efferents, and was uniquely distinguished by the absence of modulation in thalamic efferents (Fig. 2n).

### Low–high mystical experience differences in psilocybin-modulated effective connectivity

To assess whether the influence of structural connectivity on effective connectivity differed between those that reported (relatively) high or low subjective effects under psilocybin, we repeated the procedures utilized to assess baseline–psilocybin effective connectivity changes, focusing instead on the psilocybin session, and separating subjects according to whether their MEQ scores fell above (*n* = 31, 12 female, mean age = 39.61, s.d. = 9.72) or below (*n* = 30, 16 female, mean age = 34.90, s.d. = 11.67) the fiftieth percentile. Fig. 3a–d shows that across contexts, the evidence-weighted prior-variance transformations did not differ markedly between low and high MEQ groups: there was only modest evidence for a decreased intercept at high MEQ scores in the meditation context (posterior probability = 0.72, Fig. 3b). Again, applying these evidence-weighted prior-variance transformation led to a marked increase in evidence at both the group (Fig. 3e) and subject level (SI Results, Fig. S3).

**Fig. 3.**
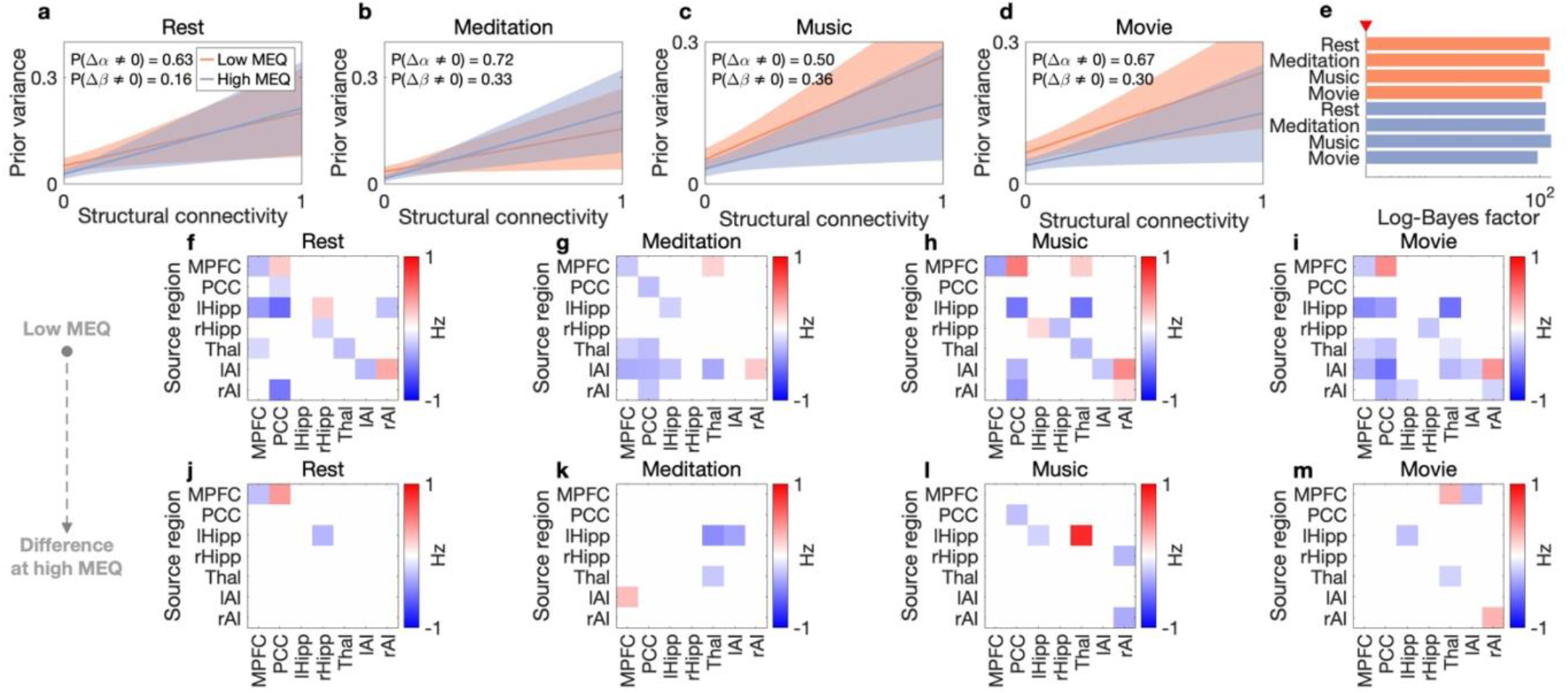
Low and high mystical experience subgroups show structurally informed effective connectivity differences. (**a–d**) Evidence-weighted prior-variance transformations derived from high (blue) and low (orange) mystical experience questionnaire (MEQ) subgroups (median split). Shading denotes standard error envelope. Modest evidence that intercepts (*α*) are lower for high-MEQ subjects across contexts, indicating lower baseline variability in effective connectivity (with no clear change in slope, *β*). Two-tailed posterior probability of low–high MEQ parameter differences reported in top left of each panel. (**e**) Log-Bayes factors showing increase in group-level evidence seen when each hierarchical empirical Bayes model was inverted under its evidence-weighted prior-variance transformation (orange: low MEQ; blue: high MEQ). Note red triangle indicates log-Bayes factor of 3 (x-axis is log scaled). Subject-level log-Bayes factors not shown (SI Results, Fig. S3). (**f–i**) Effective connectivity in the low MEQ group, resembling the group-level psilocybin profiles in Fig. 2. Self-connections (diagonal entries) are scaled relative to a fixed inhibitory prior (−0.5 Hz) via a log-normal transformation; values reflect relative changes, not absolute Hz. (**j–m**) Differential connectivity in the high MEQ group, most notably in the meditation condition (**k**), where efferent connectivity from bilateral hippocampus to anterior insula and afferent input to PCC were modulated (summed matrices depicting the full psilocybin profiles, SI Results, Fig. S4).

Leveraging structure-based priors furnished by the MEQ group prior-variance transformations (Fig. 3a–d), we inverted hierarchical empirical Bayes models to quantify low–high MEQ-related differences in effective connectivity (Methods). The low-MEQ group’s profiles (Fig. 3f–i) were largely consistent with the psilocybin-modulated connectivity patterns identified for the full cohort (Fig. 2; SI Results, Fig. S2). Across contexts, differences at high MEQ scores (Fig. 3j–m; SI Results, Fig. S4) were most pronounced for efferents from the left hippocampus. Notably, effective connectivity from the left hippocampus to thalamus was downregulated in the guided meditation context and upregulated during music listening (Fig. 3k–l).

### Predictive validity of psilocybin-modulated effective connectivity of the left hippocampus

In light of the consistencies and differences observed across prior analyses, we next used a hierarchical empirical Bayes model that treated MEQ scores as a continuous covariate and assessed MEQ-dependent changes in psilocybin-modulated effective connectivity, performing leave-one-out cross-validation on model variants that focused on left-hippocampal efferents (Methods). Fig. 4a–d shows the MEQ-dependent component of psilocybin-modulated effective connectivity (mean connectivity under psilocybin, independent of MEQ, is reported in SI Results, Fig. S5). Although the strength modulation was attenuated relative to the split-group analyses, the pattern of MEQ-related modulation of left-hippocampal efferents overlapped with that shown in Fig. 3j–m.

**Fig. 4.**
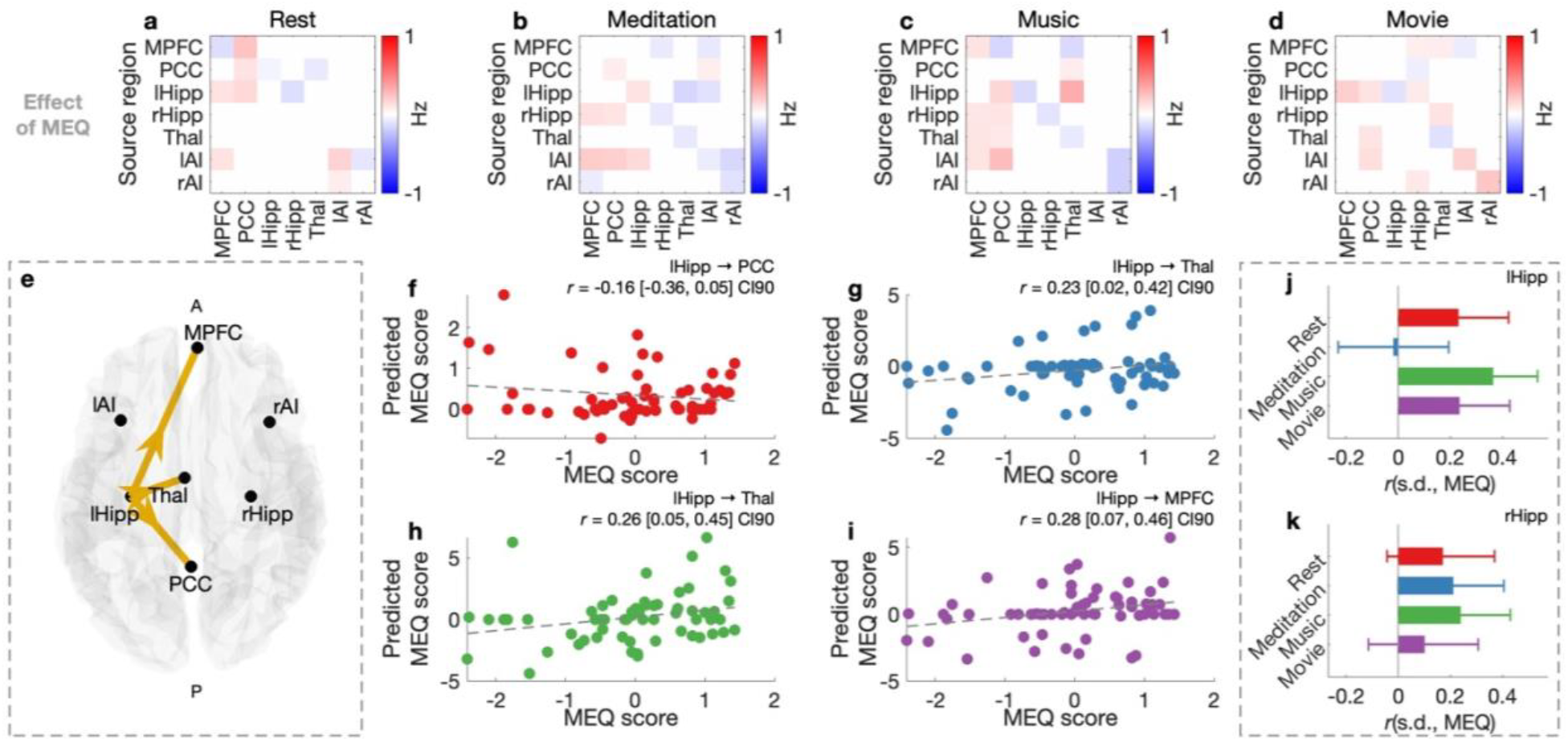
Structurally informed psilocybin-modulated effective connectivity predicts mystical experience. (**a–d**) Group-level changes in effective connectivity associated with mystical experience questionnaire (MEQ) scores during rest (**a**), guided meditation (**b**), music listening (**c**), and movie viewing (**d**). (**e**) Graphical depiction of the three effective connections—efferents from the left hippocampus (lHipp)— considered in leave-one-out cross-validation analyses: lHipp to posterior cingulate cortex (PCC), lHipp to thalamus (Thal), and lHipp to medial prefrontal cortex (MPFC). These connections represent the strongest changes to lHipp efferents shown in initial panels. (**f–i**) Observed versus predicted MEQ scores from leave-one-out cross-validation for lHipp to PCC connection during rest (**f**), lHipp to Thal connection during guided meditation (**h**) and music listening (**i**), and lHipp to MPFC connection during movie viewing (**g**). Pearson correlation and 90% confidence intervals are reported for each comparison. (**j, k**), Correlation between the standard deviation of regional blood-oxygen-level-dependent (BOLD) signals and MEQ scores for lHipp (**j**) right hippocampus (rHipp) (**k**) across contexts (under psilocybin). Error bars indicate 90% confidence intervals.

Focusing on the strongest MEQ-modulated left-hippocampal efferents (Fig. 3a–d)—namely, the connection to the PCC (resting state), thalamus (guided meditation and music listening), and MPFC (movie viewing; arrows, Fig. 4e)—we conducted leave-one-out cross-validation analyses following an approach described by Zeidman and colleagues^28^, which emphasizes assessment of single connections of interest (avoids overparameterization). Briefly, for each context and connection of interest, and for each of the *s* = 1, …, *S* iterations (where *S* is the number of subjects), a hierarchical empirical Bayes model treating MEQ scores as a covariate was inverted using the data from all but the *s*-th (left-out) subject (Methods). The left-out subject’s MEQ score was then predicted using the group-level model and that subject’s data (first-level parameter).

Leave-one-out cross-validation revealed modest predictive accuracy (correlation range = −0.16–0.28, Fig. 4f– i; visualizations with posterior predictive covariance, SI Results, Fig. S6). The correlation between observed and predicted MEQ scores was negative for the left-hippocampus–PCC connection during rest (correlation reported with 90% confidence interval, consistent with posterior-probability thresholding, Fig. 4f), but positive for the left-hippocampus–thalamus and left-hippocampus–MPFC connections in the other (naturalistic imaging) contexts (Fig. 4g–i). As a benchmark for minimal, non-mechanistic information, we examined whether regional blood-oxygen-level-dependent (BOLD) signal variability (standard deviation) in the hippocampus—a simple local measure linked to psychedelic-modulated signal dispersion^7^—correlated with MEQ scores. Across contexts, several modest correlations (range = −0.02–0.36) were observed for both hemispheres (Fig. 4j–k), with larger effects for the left hippocampus. This convergence between structurally informed effective connectivity and model-free signal-variability measures is consistent with the view that psilocybin’s subjective effects may be preferentially associated with left-lateralized hippocampal dynamics.

## Discussion

We combined tractography-derived structural connectivity with dynamic causal modeling in a hierarchical empirical Bayes framework to assess how psilocybin alters effective connectivity across four experiential contexts (rest, guided meditation, music listening, and movie viewing). Evidence-weighted, structure-based priors improved model evidence, and the clearest psilocybin-modulated mystical experience-related effects centered on left-hippocampal efferents—most notably the left-hippocampus-to-thalamus pathway, whose MEQ association flipped sign across contexts (downregulation during guided meditation and upregulation during music listening). Leave-one-out cross-validation analyses yielded modest predictive accuracy for MEQ scores from single left-hippocampal efferents, and model-free signal variability (regional BOLD standard deviation) in left hippocampus revealed a convergent MEQ scaling.

Our findings extend prior work showing that psychedelic states are accompanied by increased variance or entropy in neural signals^3,5,16^, often interpreted in terms of a loosening of structural constraints on brain dynamics^6,7^. We observed a similar broadening in the domain of effective connectivity, but crucially, this manifested as an intercept shift in the movie viewing condition only—reflecting a change in the overall magnitude of variance—rather than a change in the slope that indexes the scaling of variance with structural connectivity. This pattern suggests that psilocybin may, in certain contexts, increase the noisiness of inter-regional interactions without altering the degree to which those interactions are constrained by structural connectivity. One possibility is that these changes arise from local, geometry- or grey matter-mediated processes rather than long-range white-matter-tract decoupling^29^, though this remains speculative. Here, it is also important to note that the variance of effective connectivity is nonlinearly related to the variance of both neural signals and functional connectivity (SI Results, Fig. S7).

We build on prior work linking classic psychedelics to hippocampal engagement (and disrupted hippocampal—default-mode coupling) in humans by moving from undirected functional connectivity to directed effective connectivity^3,30^, isolating left-hippocampal efferents as a plausible mechanism for psilocybin-related mystical experience intensity and showing that the sign of the MEQ–connectivity relationship is context dependent (in guided meditation and music listening context). Interpreted against the thalamus’ role as a task-dependent gate^31^, downregulation (increasing inhibition) of left-hippocampus-to-thalamus connectivity during meditation is compatible with reduced mnemonic or associative demands when attention turns inward^32^, whereas its upregulation (decreasing inhibition) during music listening is consistent with recruitment of autobiographical or associative processes^12^. Because the magnitude and direction of this regulation covaried with MEQ scores, one parsimonious view is that higher MEQ scores reflect greater state-transition capacity—lower control energy required to reach the context-appropriate regime under psilocybin— though individual initial states (initial distance from those regimes) likely contribute as well^33^. On this view: although psychedelics do appear to flatten the brain’s control-energy landscape^34^, a direct link between control-energy metrics and subjective intensity remains to be demonstrated.

This study represents one of the first applications of the hierarchical empirical Bayes model outside the initial methodological work in which it was introduced^17^, and—to our knowledge—the first application of a structurally informed model of directed connectivity in the context of psychedelic research. By embedding structure-based priors into an effective connectivity model, this approach allowed us to assess both psilocybin- and MEQ-related changes in the evidence-weighted mapping from structural connectivity to the prior variance of effective-connectivity parameters, and to impose biophysically grounded regularization on the inference of effective connectivity. Beyond the present application, this framework offers a means of evaluating how anatomical constraints shape and reshape neurodynamic states across pharmacological and behavioral manipulations, particularly in conditions where structure–function coupling differences have been identified using simple correlation-based measures^29^.

Several limitations motivate future directions. First, with our previous methodological work focused on large-scale, data-derived networks^17^, here, we deliberately examined a small network of seven regions implicated in psychedelic action, in line with (archetypal) hypothesis-driven DCM analyses^35^. This restricted scope may limit the generalizability of our findings, and future work should extend these analyses to the whole brain. Second, modeling the thalamus and hippocampus as single nodes sacrifices subnuclear and subfield specificity. Future work can hope to exploit high-field imaging and to localize pathways (for example, the subiculum to anterior thalamus pathway^36^) implicated by our results. Finally, while our intercept-based finding invites the hypothesis that psilocybin’s effect on effective connectivity variance may be mediated by grey matter connectivity rather than long-range white matter pathways, this remains to be tested. In future, approaches that involve the eigendecomposition of cortical surface geometry could be integrated into the hierarchical empirical Bayes model to evaluate this possibility^9,37^.

In conclusion, psilocybin reorganized effective connectivity while preserving structure-based scaling, in a context-dependent manner tied to subjective intensity. Effects centered on left-hippocampal efferents, with a sign-reversing modulation of left-hippocampus-to-thalamus—downregulation in guided meditation and upregulation in music—that tracked MEQ scores. These findings show that structurally informed effective connectivity can link brain structure, directed connectivity dynamics, and experience under psychedelics, offering a framework for understanding how context shapes the impact of these compounds on the brain.

## Methods

### Data

This study utilized open-access data from PsiConnect^15,16^, which to date, is the largest single-site neuroimaging dataset of psychedelic-naive healthy adults. Sixty-one subjects with complete data—diffusion-weighted MRI and fMRI recorded during resting state, guided meditation, music listening and movie viewing—were included in our analyses (Fig. 1a–d). Data also included responses on the MEQ, the 30-item self-report questionnaire developed and validated as an index of the intensity of psilocybin-modulated altered states of consciousness^19,38^. Essential methodological details concerning fMRI and structural connectivity are summarized below. Additional information is provided in the supporting information (SI Methods), which details priors over all free parameters (Table S1). Detailed acquisition procedures are described in the dataset paper by Novelli and colleagues^15^.

#### Imaging acquisition and preprocessing

Resting-state and (naturalistic) task-based fMRI data were acquired on a Siemens 3T Magnetom Skyra scanner equipped with a 32-channel head coil and enhanced gradient system. Functional images were collected using a multi-echo, multi-band echo-planar imaging (EPI) sequence: repetition time (TR) of 910 ms, echo time (TEs) of 12.60 ms, 29.23 ms, 45.86 ms, 62.49 ms, multi-band acceleration factor of 4, field of view (FOV) of 206 mm, right–left phase encoding direction, and 3.2 mm isotropic voxels. Data were acquired during four conditions: resting state (8 min, 505 volumes), audio-guided meditation (6 min 30 s, 405 volumes), music listening (11 min 24 s, 728 volumes), and movie viewing (6 min, 372 volumes). Multi-echo fMRI data were preprocessed with fMRIPrep 22.0.2, including standard structural corrections, surface reconstruction, and spatial normalization to Montreal Neurological Institute (MNI) space MNI152NLin2009cAsym^39,40^. Data were denoised in the Statistical Parametric Mapping (SPM) toolbox (version 12) via regression of white matter, cerebrospinal fluid, framewise displacement (head motion), and non-BOLD components identified via multi-echo independent components analysis (tedana 0.0.12)^41,42^. Preprocessed fMRI data were then parcellated by extracting mean BOLD time series from regions of interest (ROIs) defined in MNI space and mapped to voxel coordinates. ROIs were spherical (radius 6–8 mm) centered on MNI coordinates from prior literature (SI Methods).

Subject-level structural connectivity utilized in this study was derived from whole-brain probabilistic tractography applied to subject’s diffusion-weighted MRI acquired using an echo-planar imaging sequence (TR = 5900 ms, TE = 171 ms, 2.5 mm isotropic voxels) with two b-value shells (0 and 3,000 s/mm^2^). Following standard MRtrix3 pipelines, fiber orientation distributions were reconstructed in each subject’s native T1 space using the iFOD2 algorithm^43^, and anatomically constrained tractography (ACT) was used to dynamically seed and generate 4 million streamlines per subject^44^ (further information, SI Methods). Streamlines were parcellated using a custom atlas comprising spherical ROIs. Group-average structural connectivity was computed by re-weighting individual streamline count matrices for inter-ROI distance and size, then averaging across subjects and normalizing by the strongest (average) connection.

### Model

Effective connectivity was inferred using a hierarchical empirical Bayes framework built on the well-validated spectral DCM for resting-state fMRI^18,45,46^, and standard parametric empirical Bayes routines implemented in the SPM toolbox^24^. Full details of the generative model, prior specification, and inversion procedure— including in silico and empirical validation—are reported in our previous work and are detailed in the Supplementary Methods^17^. Here, we provide a high-level summary of the framework and highlight a few novel aspects—particular to this study—concerning the use of design matrices and structure-based priors for between-subject (within-subject) effective connectivity differences (changes).

Starting at the first (subject) level, for *s* = 1, …, *S* subjects, and each context, we consider a DCM that describes how *i* = 1, …, *n* neuronal populations influence one another and give rise to observed BOLD responses. The temporal-domain formulation assumes the following continuous-time state-space representation:

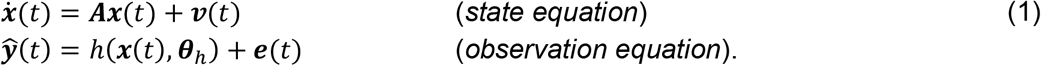

Here, ***x***(*t*) = [*x*_1_(*t*), …, *x*_*n*_(*t*)]^T^ denotes the neuronal state vector, where each scalar function *x*_*i*_(*t*) ∈ ℝ describes the ensemble (or mean-field) activity of the *i*-th population at time *t*. The transition matrix ***A*** ∈ ℝ^*n*×*n*^ encodes the intra- and inter-regional modulation of the rates of change in this ensemble activity (the effective connectivity), in its diagonal and off-diagonal elements, respectively. In the observation equation, *h* is the hemodynamic response function (HRF) with free parameters ***θ***_*h*_ (SI Methods), which maps ensemble neuronal activity to expected BOLD responses ***ŷ*** (*t*), which are compared to empirical data with parameterized residual precision (SI Methods). Both endogenous fluctuations ***v***(*t*), and observation error ***e***(*t*), are parameterized as power-law noise (SI Methods).

Using the Fourier transform, the state-space model is transformed into the spectral domain, such that it generates the expected cross-spectral density (CSD) of BOLD responses (for brevity, we forgo a detailed account of this spectral transformation and direct the reader to Novelli and colleagues for a didactic introduction to this aspect of the model^47^). Then, model inversion proceeds using variational Bayes under the Laplace approximation (VBL)^48^, which iteratively updates the sufficient statistics of (approximate) Gaussian posteriors over parameters to maximize free energy: a lower bound on the log model evidence.

With DCMs inverted for a specified network of regions, the posteriors of interest—for effective connectivity, in this context—are moved into the hierarchical empirical Bayes model. To obtain an evidence-weighted prior-variance transformation—per context and session (Fig. 2b–e), or per context and MEQ subgroup (Fig. 3a–e), for example—we proceed as follows. Let ***θ***^(1)^ be a vector stacked MAP estimates (posterior means) of interest for *S* subjects, such that 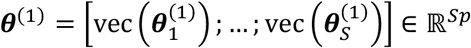. Here, 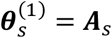 are the effective connectivity parameters for the *s*-th subject (obtained, for example, under psilocybin in the resting-state context), and *p* = *n*^2^ is the number of the *s*-th subject’s parameters considered (Fig. 1i). We can now consider a hierarchical regression formulation:

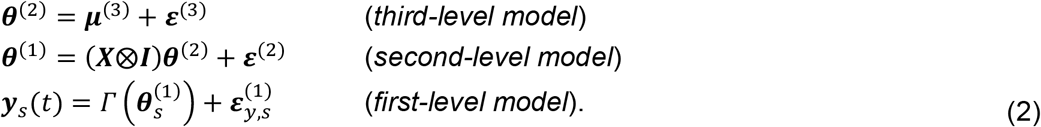

Here, at the third level, ***θ***^(2)^ ∈ ℝ^*pq*^ is the group-level effective connectivity (Fig. 1h) modeled as deviations ***ε***^(3)^~𝒩(**0, Σ**^(3)^) from prior expectations ***μ***^(3)^ (Fig. 1g), and *q* is the number of covariates (including the intercept) encoded in a second-level design matrix ***X*** ∈ ℝ^*S*×*q*^ (to obtain the prior-variance transformation, we use the intercept-only case: *q* = 1, ***X*** = **1**_*S*×1_). At the second level, the Kronecker product ***X***⨂***I*** ∈ ℝ^*Sp*×*pq*^— where ***I*** ∈ ℝ^*p*×*p*^ is the identity matrix—ensures that both ***θ***^(2)^ and the subject-specific random effects ***ε***^(2)^, are appropriately tiled across parameters in ***θ***^(1)^. Following conventions, ***ε***^(2)^~𝒩(**0, Σ**^(2)^) is parametrized in terms of one or more scaled precision components (Supplementary Methods). The final line of Eq. 3 models the observed data for each subject ***y***_*s*_(*t*), as the output of the DCM mapping *Γ* applied to subject-specific effective connectivity 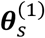 plus residual error 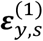. Although in this context *Γ* maps parameters to the observed CSD, we retain the time-domain representation for consistency with Eq. 1 (Fig. 1j, SI Methods).

At the third level, structural connectivity is incorporated into the prior covariance **Σ**^(3)^ via the following:

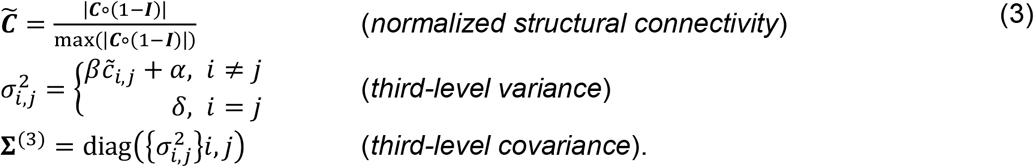

Here, 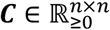 encodes structural connectivity for the *n*-region network (Fig. 1e), ∘ denotes the elementwise product, (1 − ***I***) is the complement of the *n*-dimensional identity matrix, and the normalized structural connectivity weights 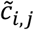 from the *i*-th row and *j*-th column are transformed to produce variance terms 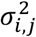 for corresponding group-level effective connections ***θ***^(2)^. The hyperparameters *α* and *β* set the baseline variance and its scaling with structural connectivity, respectively, and intraregional connections—where *i* = *j*—are set to *δ* = 1/64 (Fig. 1f).

Inverting an uninformed model—the case where *α* = 1/2 and *β* = 0—and obtaining posteriors over second-level model parameters ***θ***^(2)^, affords the opportunity to explore (reduced) structurally informed models that are nested within this (full) uninformed model by varying *α* and *β*, and obtaining updated posteriors and free energy using BMR^24^. By weighting the *k*-th set of evaluated hyperparameters 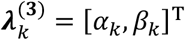, by their corresponding posterior model probabilities 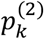, computed via a softmax transformation of the model free energies 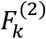, the hyperparameters of the evidence-weighted prior-variance transformation can be obtained following:

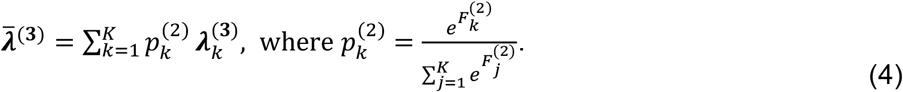

This BMA procedure serves to identify the parameters of a parsimonious transformation (avoid overfitting) in a manner that retains the influence of plausible alternatives transformations^25^. Uncertainty in 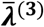 can be quantified as the standard deviation of the evidence-weighted samples 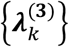 under 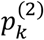, and propagated to the transformed prior variances using the delta method (furnishing standard error envelopes, Figs. 2–3).

Let us now consider the scenario where one has obtained evidence-weighted hyperparameters for two sessions (baseline and psilocybin, Fig. 2b–f) or subgroups (low and high MEQ, Fig. 3a–d) and wishes to investigate effective connectivity changes or differences using the model described in Eq. 2. In the case of splitting subjects according to their MEQ scores, we can encode the effect of being above or below the sample median with a binary indicator variable. The second-level design matrix becomes:

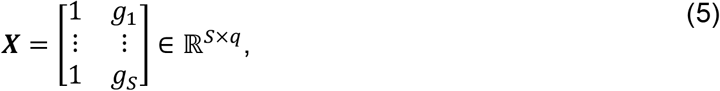

where the first column now models the mean effective connectivity for the low MEQ group, and *g*_*s*_ ∈ {0,1} is the group indicator for the *s*-th subject (*g*_*s*_ = 1 for high MEQ). Let 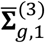 and 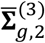 be the third-level prior covariance obtained under evidence-weighted hyperparameters for the low and high MEQ groups (in a particular context), respectively. To construct the third-level prior covariance **Σ**^(3)^ for a structurally informed model encoding low–high MEQ differences, we extract the variance terms (diagonals) from these group-specific covariance matrices and assign them to the appropriate design-matrix columns:

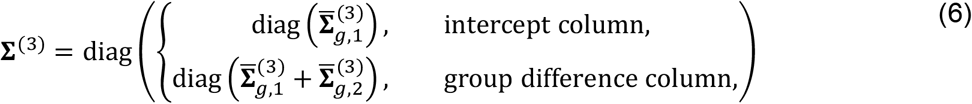

Here, diag(·) extracts the diagonal elements of a matrix into a column vector, so that **Σ**^(3)^ is diagonal in the vectorized parameter space. The intercept column encodes the low-MEQ variance, while the group-difference column encodes the sum of variances from the low and high MEQ groups. Note that the corresponding third-level prior mean ***μ***^(3)^ is constructed in an analogous manner, and that before inversion, **Σ**^(3)^ is rescaled to ensure design-scale invariance across regressors (SI Methods).

Although Eqs. 5–6 focus on between-group differences, the same procedure extends naturally to within-subject contrasts—baseline versus psilocybin, for example—by stacking parameters from both sessions into ***θ***^(1)^ and configuring the design matrix ***X*** to code for session identity in place of group membership. In both cases, the approach is identical in spirit: structure-based priors are assigned to the relevant design-matrix columns, and the model is inverted under these priors to test for the corresponding effects.

### Procedures

Focusing on a network of seven brain regions chosen *a priori* for mechanistic relevance to psychedelic action (Fig. 2a), for each session (baseline and psilocybin) context (resting-state, guided meditation, music listening and movie viewing), spectral DCMs were inverted separately for each subject to obtain posterior estimates of effective connectivity. These subject-level posteriors were then taken to a hierarchical empirical Bayes model with an intercept-only design matrix at the second level, and uninformed priors at the third level. After inverting this model, we utilized grid search and BMR to evaluate the evidence associated with different reduced (structurally informed) group-level models, varying the parametrization of the prior-variance transformation (Eq. 3). In these analyses, *α* and *β* were sampled at 100 equidistant points across [0, 1/2]. Valid parameter combinations, satisfying *α* + *β* ∈ [10^−5^, 1/2] were retained, yielding a triangular sampling space with *K* = 5049 parameter regimes.

After obtaining the evidence-weighted hyperparameters using BMA (Eq. 4), uncertainty in these parameters was quantified as the evidence-weighted standard deviation and propagated to the transformed prior variances using the delta method (Fig. 2b–e). This uncertainty quantification also afforded us the ability to assess the posterior probability of baseline–psilocybin differences in evidence-weighted hyperparameters (top left, Fig. 2b–e). Reinverting these session- and context-specific hierarchical empirical Bayes models *de novo*, under their respective evidence-weighted prior-variance transformations, permitted quantification of the increase in evidence afforded by structure-based priors in terms of the log-Bayes factor (difference in free energies for the uniformed and structurally informed models, Fig. 2f, Fig. S1). Finally, for within-subject contrasts (Fig. 2g–n), parameters from both sessions were entered into hierarchical empirical Bayes models with an appropriately specified second-level design matrix and third-level prior (Eqs. 5–6).

These procedures were repeated for each MEQ subgroup and context within the psilocybin session, yielding evidence-weighted prior variance transformations with estimated uncertainty (Fig. 3a–d), the increase in evidence afforded by structure-based priors (Fig. 3e, Fig. S3), and between-subject contrasts (Fig. 3f–m). Prior to leave-one-out cross-validation analyses, we also estimated the MEQ effect dimensionally (Fig. 4a– d): MEQ scores were treated as a continuous covariate, and the group-level priors on the MEQ coefficients were set as a scaled copy of the intercept priors derived from the prior-variance mapping (SI Methods). Across results, we used a two-tailed 90% posterior-probability threshold (per-parameter false-sign ≤ 10%).

For leave-one-out cross-validation analyses (psilocybin session), each fold focused on a single pre-specified left-hippocampal efferent (Fig. 4e). The session- and context-specific prior-variance transformation was held fixed across folds, such that structure-based priors exerted local (edgewise) shrinkage only, rather than a global matrix-wise influence. Nevertheless, the degree of connection-specific regularization was retained from the earlier steps. Prediction accuracy was summarized with Pearson’s correlation and two-sided 90% Fisher-z confidence intervals (chosen for consistency with posterior-probability thresholding).

## Data availability

The PsiConnect dataset used in this study is undergoing final curation and is currently under embargo. Upon publication of the associated data descriptor^15^, the dataset will be released on OpenNeuro (accession ds006110). During peer review, de-identified derivatives sufficient to reproduce the analyses will be made available to the handling editors and reviewers via a private repository.

## Code availability

Analyses utilizing the hierarchical empirical Bayes model were conducted using custom MATLAB scripts developed for this study, building on core routines from the SPM toolbox. Diffusion MRI preprocessing and tractography were performed using MRtrix3, FSL, and ANTs, following the pipeline described in the SI Methods, with custom scripts for structural connectivity matrix construction and parcel registration. Code and scripts, along with derived data sufficient to reproduce the key analyses and figures, will be made publicly available via GitHub upon publication.

## Acknowledgements

M.D.G. and T.B. are supported by Australian Government Research Training Program Scholarships. M.D.G., L.N. and A.R. are funded by the Australian Research Council (ref. DP200100757). A.R. is also funded by Australian National Health and Medical Research Council Investigator Grant (ref. 1194910). A.R. is affiliated with The Wellcome Centre for Human Neuroimaging supported by core funding from Wellcome (203147/Z/16/Z). A.R. is a CIFAR Azrieli Global Scholar in the Brain, Mind & Consciousness Programme.

## Author contributions

M.D.G., T.B. and A.R. conceived the study. M.D.G. and T.B. developed the methodology, implemented the software, performed the formal analyses and investigations, curated the data, and contributed to visualization and project administration. M.D.G. additionally performed validation, drafted the original manuscript and secured funding. L.N. contributed to data curation and supervision. All authors contributed to the review and editing of the manuscript. A.R. also provided supervision and funding acquisition.

