## Supporting information for "Context-dependent structurally informed effective connectivity under psilocybin"

#### Overview
This supporting information (SI) document provides additional analyses, figures, and methodological details that support the findings reported in the main text. Elements appear in the order in which they are referenced in the main text. A complete list of contents is provided in the table below for ease of navigation.

### SI Methods

#### Completing the hierarchical empirical Bayes model

##### Dynamic causal modeling at the first level

To complete the specification of the hierarchical empirical Bayes model (Methods, Eqs. 1–2), we detail the form of the HRF, endogenous fluctuations, observation error, the construction of the empirical CSD matrix, the parametrization of residual error (in the context of VBL) and the second-level prior covariance for random effects (RFX). In Eq. 1, $h(\boldsymbol{x}(t), \boldsymbol{\theta}_h)$ represents the well-known 'Balloon' HRF that yields expected BOLD responses from regional neuronal states $x_i(t)$ as follows[1]:

$$\begin{aligned}
\dot{s}_i(t) &= x_i(t) - k_h \cdot s_i(t) - \gamma_h \cdot (f_i(t) - 1), && \textit{(vasodilatory signal)}\\
\dot{f}_i(t) &= s_i(t), && \textit{(blood flow induction)}\\
\dot{b}_i(t) &= \frac{1}{\tau_i} \cdot f_i(t) - b_i(t)^{1/\alpha_h}, && \textit{(blood volume)}\\
\dot{q}_i(t) &= \frac{1}{\tau_i} \cdot \left( \frac{f_i(t) \cdot \left(1 - (1 - E_0)^{1/f_i(t)}\right)}{E_0} - b_i(t) \cdot q_i(t)^{1/\alpha_h - 1} \right), && \textit{(deoxyhemoglobin content)}\\
\hat{y}_i(t) &= V_0 \cdot \left[ k_1 \left(1 - q_i(t)\right) + k_2 \left(1 - \frac{q_i(t)}{b_i(t)}\right) + k_3 \left(1 - b_i(t)\right) \right]. && \textit{(BOLD responses)}
\end{aligned} \quad \text{(S1)}$$

Here, neuronal activity $x_i(t)$ drives a vasodilatory signal $s_i(t)$, which increases blood flow $f_i(t)$, expands blood volume $b_i(t)$ via Grubb's law, alters deoxyhemoglobin content $q_i(t)$ through oxygen extraction, and together these variables determine the expected BOLD response $\hat{y}_i(t)$. The vasodilatory signal decays at rate $k_h$, the feedback of blood flow is regulated by $\gamma_h = 0.32$, $\boldsymbol{\tau} = [\tau_1, \ldots, \tau_n]$ are region-specific transit times, $\alpha_h = 0.32$ is Grubb's (vessel stiffness) exponent, and $E_0 = 0.4$ and $V_0 = 4$ represent the resting oxygen extraction fraction and resting venous blood volume fraction, respectively. In the last line of Eq. S1, coefficients $k_1$, $k_2$, and $k_3$ represent the contributions of the intra- and extra-vascular compartments to the BOLD response:

$$k_1 = 4.3 \cdot v_0 \cdot E_0 \cdot TE, \qquad k_2 = \epsilon_h \cdot r_0 \cdot E_0 \cdot TE, \qquad k_3 = 1 - \epsilon_h, \quad \text{(S2)}$$

where $v_0 = 40.3$ is the frequency offset at the outer surface of magnetized vessels, $TE = 0.04$ is the echo time, $r_0 = 25$ is the slope of the intravascular relaxation rate, and $\epsilon_h$ is the ratio of intra- to extra-vascular signal contributions. The set of parameters that were free to vary is given by $\boldsymbol{\theta}_h = \{k_h, \boldsymbol{\tau}, \epsilon_h\}$.

Using the Fourier transform, $\mathcal{F}$, these state-space equations (Eq. 1 and Eq. S1) are transformed into the spectral domain, such that they generate the expected CSD of BOLD responses $\widehat{\boldsymbol{G}}_y(\omega) = \left[\mathcal{F}\{\hat{\boldsymbol{y}}(t)\}\mathcal{F}\{\hat{\boldsymbol{y}}(t)\}^\dagger\right]$, where † denotes the conjugate transpose, and the right-hand side is implicitly a function of angular frequency $\omega$ via the Fourier transform. Putting this all together, the spectral equivalent of Eq. 1 reads:

$$\widehat{\boldsymbol{G}}_y(\omega) = \boldsymbol{H}(\omega)(i\omega\boldsymbol{I} - \boldsymbol{A})^{-1}\boldsymbol{G}_v(\omega)(-i\omega\boldsymbol{I} - \boldsymbol{A}^T)^{-1}\boldsymbol{H}(\omega)^\dagger + \boldsymbol{G}_e(\omega), \quad \text{(S3)}$$

where $\boldsymbol{H}(\omega)$ is the Fourier transform of the HRF. Here, notably, the latent neuronal state in the frequency domain, $\boldsymbol{X}(\omega)$, has been factored out via the substitution $\boldsymbol{X}(\omega)\boldsymbol{X}(\omega)^\dagger = (i\omega\boldsymbol{I} - \boldsymbol{A})^{-1}\boldsymbol{G}_v(\omega)(-i\omega\boldsymbol{I} - \boldsymbol{A}^T)^{-1}$. The model of endogenous fluctuations $\boldsymbol{v}(t) \to \boldsymbol{G}_v(\omega) \in \mathbb{R}^{n\times n}$, and observation error $\boldsymbol{e}(t) \to \boldsymbol{G}_e(\omega) \in \mathbb{R}^{n\times n}$ (Eq. S3). Both models take the form of a diagonal matrix-valued function of angular frequency $\omega$, whose entries—$g_{v,i,i}(\omega)$ and $g_{e,i,i}(\omega)$, respectively—encode an auto-spectrum or power spectral density (PSD) model of region-specific endogenous fluctuations and observation error:

$$g_{v,i,i}(\omega) = \alpha_v \omega^{-\beta_v}, \qquad g_{e,i,i}(\omega) = (\gamma_e \cdot \alpha_{e,i}) \cdot \frac{\omega^{-\beta_e/2}}{\sum_\omega \omega^{-\beta_e/2}}. \quad \text{(S4)}$$

Here, the PSD model for endogenous fluctuations is parameterized by a global amplitude $\alpha_v$ and spectral exponent $\beta_{v,i}$, whereas the PSD model for observation error is controlled by a global spectral exponent $\beta_e$, and region-specific amplitude parameter $\alpha_{e,i}$ normalized by a global factor $\gamma_e$.

To construct the empirical CSD matrix $\boldsymbol{G}_y(\omega)$ for model inversion, we fit a multivariate autoregressive model of order $L = 8$ to observed BOLD time series using the variational Bayesian procedure described by Penny and Roberts[2]. This procedure yields a set of autoregressive coefficient matrices $\{\boldsymbol{W}_k \in \mathbb{R}^{n \times n}\}_{k=1}^{L}$ and a residual covariance matrix $\boldsymbol{\Sigma}_w \in \mathbb{R}^{n \times n}$. The standard spectral factorization then produces $\boldsymbol{G}_y(\omega)$ as:

$$\boldsymbol{G}_y(\omega) = \boldsymbol{\mathcal{H}}(\omega)\boldsymbol{\Sigma}_w\boldsymbol{\mathcal{H}}(\omega)^{\dagger}, \text{ where } \boldsymbol{\mathcal{H}}(\omega) = \left(\boldsymbol{I} + \textstyle\sum_{k=1}^{L} \boldsymbol{W}_k\, e^{-i\omega k}\right)^{-1}, \tag{S5}$$

Consistent with prior related work, $\boldsymbol{G}_y(\omega)$ is estimated at $n_f = 32$ linearly spaced frequencies between $1/128$ Hz and the Nyquist frequency: $1/2 \cdot TR$, where $TR$ is the fMRI repetition time (sampling interval)[3].

In the context of model inversion using VBL, the expected log-likelihood quantifies the fit between the expected and observed CSDs under the approximate posterior. This is expressed as:

$$\text{vec}\left(\boldsymbol{G}_y(\omega)\right) = \text{vec}\left(\widehat{\boldsymbol{G}}_y(\omega)\right) + \boldsymbol{\epsilon}_y \tag{S6}$$

Here, both real and imaginary parts are stacked, and the residual error $\boldsymbol{\epsilon}_y \sim \mathcal{N}(\boldsymbol{0}, \boldsymbol{\Sigma}_y)$ is parameterized in terms of its precision. Per the implementation utilized here, the residual error precision $\boldsymbol{\Pi}_y = \boldsymbol{\Sigma}_y^{-1}$, is expressed as a weighted sum of $n^2$ precision components:

$$\boldsymbol{\Pi}_y = \textstyle\sum_{i=1}^{{n_r}^2} e^{\lambda_{y,i}}\, \boldsymbol{Q}_{y,i}, \tag{S7}$$

where $\boldsymbol{Q}_{y,i} \in \mathbb{R}^{(n^2 \cdot n_f) \times (n^2 \cdot n)}$ is a sparse, block-diagonal matrix with an $n_f$-dimensional identity sub-block targeting the $i$-th region pair (including self-connections). Here, the precision weights $\lambda_{y,i}$ are drawn from a hyperprior $\lambda_{y,i} \sim \mathcal{N}(8, 1/128)$, where these settings were determined elsewhere[4].

#### Random effects modelling at the second level

In this study, we permitted distinct connection-wise variances per covariate by using a block-diagonal prior covariance (Methods, Eq. 6). Prior to inversion, to ensure design-scale invariance we apply a column-wise normalization to the third-level prior covariance $\boldsymbol{\Sigma}^{(3)}$:

$$\widetilde{\boldsymbol{\Sigma}}^{(3)} = (\boldsymbol{D} \otimes \boldsymbol{I})\boldsymbol{\Sigma}^{(3)}(\boldsymbol{D} \otimes \boldsymbol{I}), \qquad \boldsymbol{D} = \text{diag}\left(\text{diag}\left(\left(\boldsymbol{X}^{\mathrm{T}}\boldsymbol{X}\right)^{-1/2}\right)\right), \tag{S8}$$

where $\boldsymbol{I}$ is the $p$-dimensional identity matrix (and in all our analyses $p = n^2$). Note that because $\boldsymbol{D}$ depends only on column norms, the procedure (and notation) is agnostic to whether $\boldsymbol{X}$ encodes between-group contrasts, within-subject changes or continuous covariates[5].

Finally, in the group-level model (Methods, Eq. 2), $\boldsymbol{\varepsilon}^{(2)} \sim \mathcal{N}(\boldsymbol{0}, \boldsymbol{\Sigma}^{(2)})$ are parametrized in terms of one or more scaled precision components. In all analyses assessing structure-based priors in intercept-only models ($\boldsymbol{X} = \boldsymbol{1}_S$), we used a single scaled precision component:

$$\boldsymbol{\Sigma}^{(2)^{-1}} = \boldsymbol{\Pi}^{(2)} = \boldsymbol{I} \otimes \left(\boldsymbol{Q}_0^{(2)} + e^{-\gamma^{(2)}}\boldsymbol{Q}_1^{(2)}\right). \tag{S9}$$

where $\boldsymbol{I}$ is an $S$-dimensional identity matrix, $\boldsymbol{Q}_0^{(2)} = e^{-8}\widetilde{\boldsymbol{\Sigma}}^{(3)^{-1}}$ is the lower-bound precision, $\boldsymbol{Q}_1^{(2)} = \widetilde{\boldsymbol{\Sigma}}^{(3)^{-1}}$, and the log-precision scale parameter $\gamma^{(2)}$ is inferred from the data (priors, Table S1). By contrast, in analyses of between- and within-group contrasts, we increased the flexibility by allowing parameter-specific shrinkage. Namely, we used an edgewise mixture of scaled precision components:

$$\boldsymbol{\Sigma}^{(2)^{-1}} = \boldsymbol{\Pi}^{(2)} = \boldsymbol{I} \otimes \left( \boldsymbol{Q}_0^{(2)} + \sum_{j=1}^{p} e^{-\gamma_j^{(2)}} \boldsymbol{E}_j \boldsymbol{Q}_1^{(2)} \boldsymbol{E}_j \right). \tag{S10}$$

Here, $\gamma_j^{(2)}$ is the log-precision scale of the $j$-th parameter, and the matrix $\boldsymbol{E}_j$ has zero for all entries except for $(j,j) = 1$. Elements $\boldsymbol{I}$, $\boldsymbol{Q}_0^{(2)} = e^{-8}\widetilde{\boldsymbol{\Sigma}}^{(3)^{-1}}$ and $\boldsymbol{Q}_1^{(2)} = \widetilde{\boldsymbol{\Sigma}}^{(3)^{-1}}$ are specified per Eq. S9. Priors for all free parameters are reported in Table S1.

| Parameter | Description | Prior mean | Prior variance |
|---|---|---|---|
| $\ln\left(-2 \cdot a_{s,i,i}^{(1)}\right)$ | Intra-regional effective connectivity | $-1/2$ | $1/64$ |
| $a_{s,i,j}^{(1)}$ | Inter-regional effective connectivity | $1/128$ | $1/2$ |
| $\ln \alpha_v$ | Global amplitude of endogenous fluctuations | $0$ | $1/64$ |
| $\ln \beta_v$ | Global spectral exponent of endogenous fluctuations | $0$ | $1/64$ |
| $\ln \alpha_{e,i}$ | Regional amplitude of observation noise | $0$ | $1/64$ |
| $\ln \beta_e$ | Global spectral exponent of observation error | $0$ | $1/64$ |
| $\ln \gamma_e$ | Normalization factor for observation error | $0$ | $1/64$ |
| $\ln k_h$ | Decay rate of the vasodilatory signal | $0$ | $1/256$ |
| $\ln \epsilon_h$ | Ratio of intra- to extra-vascular signal contributions | $0$ | $1/256$ |
| $\ln \tau_i$ | Transit times | $0$ | $1/256$ |
| $\ln\left(-2 \cdot a_{i,i}^{(2)}\right)$ | Second-level intra-regional effective connectivity | $-1/2$ | $1/64$ |
| $a_{i,j}^{(2)}$ | Second-level inter-regional effective connectivity | $1/128$ | Eq. 3 |
| $\gamma^{(2)}$ | Second-level precision scale parameter(s) | $0$ | $1/16$ |

**Table S1.** Prior mean and variance for free parameters utilized in the hierarchical empirical Bayes model. Notation is consistent with that utilized in the main text. If the model is structurally informed, the second-level inter-regional effective connectivity is a function of normalized structural connectivity, and the hyperparameters governing the prior-variance transformation (Methods, Eq. 3).

#### Data processing

Selection and extraction of regions of interest

The selection of ROIs and their size was based on previous research[6–8]. ROIs and MNI coordinates were MPFC [3, 54, -2], PCC [0, -52, 26], hippocampus (left [-29, -18, -16], right [29, -18, -16]), thalamus [-3, -9, 9], and AI (left [-34, 19, 0], right [38, 18, 0]). Each ROI time series was the mean of the voxels' activity within an 8-mm sphere for MPFC and PCC and a 6-mm sphere for all other regions.

Diffusion-weighted MRI preprocessing

Diffusion-weighted MRI data were preprocessed using MRtrix3 (version 3.0.3)[9], FSL (version 6.0.4)[10] and ANTs (version 2.4.3)[11]. Raw data were converted to MRtrix3 format and T1 images underwent reorientation, FOV cropping, and brain extraction (fractional intensity threshold = 0.3).

Preprocessing comprised sequential denoising using random matrix theory[12], Gibbs ringing artifact removal[13] and motion and distortion correction. Motion, eddy current, and susceptibility distortion correction was performed with slice-to-volume correction and outlier replacement[14–16]. Brain masks were generated (fractional intensity threshold = 0.2)[17] followed by bias field correction using the N4 algorithm[18]. Following preprocessing, systematic quality assurance (QA) was performed by two independent researchers (M.D.G and T.B). QA included assessment of eddy current outlier detection in addition to visual inspection of distortion correction and brain mask accuracy. Brain were extraction parameters iteratively adjusted until inter-rater agreement was achieved.

White matter fiber orientation distributions were estimated using multi-shell multi-tissue constrained spherical deconvolution with the dhollander algorithm for response function estimation[19,20]. Multi-tissue intensity normalization was applied across white matter, gray matter, and cerebrospinal fluid following three-tissue CSD modelling[19,21]. Anatomically-constrained tractography was performed using the iFOD2 algorithm, generating 4 million streamlines from gray-white matter interface seeds[22,23]. Five-tissue-type segmentation masks were created from T1 images using automated segmentation[24,25]. Streamline weights were weighted using SIFT2 to ensure biologically plausible tract densities[26].

Whole-brain structural connectivity matrices were constructed using a custom parcellation comprising ROIs. Atlas registration to individual diffusion space employed a two-stage approach: brain-extracted T1 images were registered to MNI152 space using ANTs symmetric normalization (antsRegistrationSyN.sh, antsApplyTransforms), followed by transformation to native DWI space via linear registration (flirt) in FSL. Connectivity matrices were generated using streamline count with SIFT2 weighting (tck2connectome). Parcellation preparation utilized label conversion (labelconvert) for standardized region-of-interest definition.

Subject-level left–right asymmetry of hippocampal structural connectivity

With reference to the structural connectivity shown in Fig. 2a: across 61 subjects (after distance and node-size correction), the strength of left-hippocampal connections exceeded their right-hippocampal counterparts most clearly for thalamus (54/61, 88.5%) and left AI (49/61, 80.3%), but rarely for PCC (22/61, 36.1%), MPFC (6/61, 9.8%), or right AI (4/61, 6.6%). Moreover, left-hippocampus-to-thalamus structural connectivity was the single strongest connection in the network in 50/61 (82.0%), with other left-hippocampal connections occasionally topping the network.

#### SI Results

**Subject-level evidence gains and effective-connectivity profile for psilocybin session**

To complement the results in Fig. 2f, we quantify how much structure-based priors improve model evidence per subject and per context. Fig. S1 shows, for each subject and for both sessions, the log-Bayes factor comparing the hierarchical empirical Bayes model inverted with the evidence-weighted prior-variance transformation against the uninformed model. In addition, we report the full psilocybin-session effective-connectivity profiles, obtained as the sum of the baseline matrices (Fig. 2g–j) and their corresponding psilocybin-induced changes (Fig. 2k–n). Fig. S2 displays these unthresholded psilocybin profiles with self-connections (log-normal parameters) converted back to the linear scale (Hz).

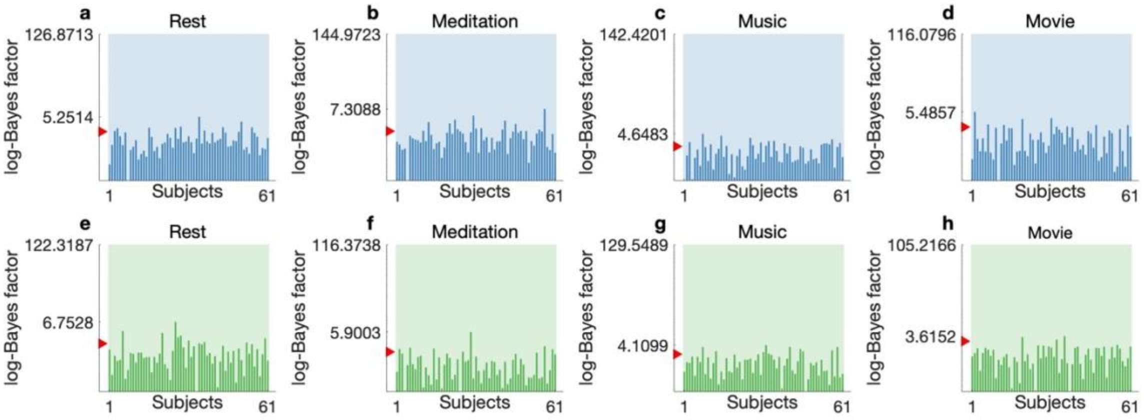

**Fig. S1 | Subject-level evidence gain from structure-based priors in whole cohort analyses.** Panels show log-Bayes factors for each context: resting state (**a, e**), guided meditation (**b, f**), music listening (**c, g**), and movie viewing (**d, h**). Results shown for baseline (blue) (**a–d**) and psilocybin (green) (**e–f**) sessions. Opaque bars are per-subject log-Bayes factors comparing the hierarchical empirical Bayes model inverted with the evidence-weighted prior-variance transformation against the uninformed model. The semi-transparent background bar (spanning the panel width) is the group-level log-Bayes factors for the same comparison. The y-axis is on a log scale; tick labels mark the largest subject-level log-Bayes factors (lower tick) and the group-level log-Bayes factors (upper tick). The red triangle indicates log-Bayes factors of 3, a conventional benchmark for strong evidence. Higher values indicate stronger support for structure-based priors.

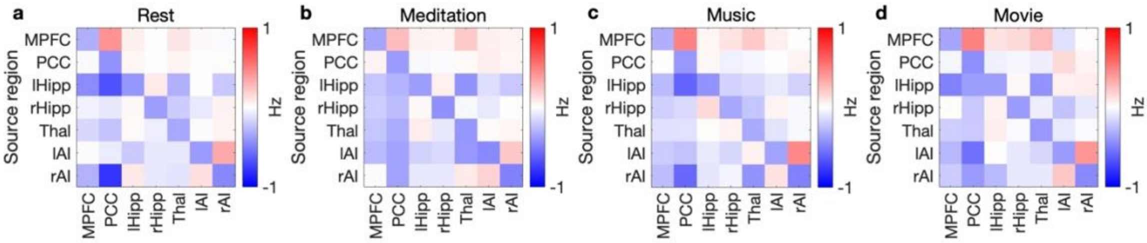

**Fig. S2 | Full psilocybin effective-connectivity profiles (unthresholded).** For each context—resting state (**a**), guided meditation (**b**), music listening (**c**), and movie viewing (**d**)—matrices depict the sum of baseline group-level effective connectivity (Fig. 2g–j) and the psilocybin-induced change (Fig. 2k–n): the full psilocybin profile. Off-diagonal entries are directed inter-regional influences (Hz). Diagonal (self-connection) parameters have been converted from the log-normal parameterization back to the linear scale.

#### Subject-level evidence gains and effective-connectivity profile for high mystical experiences

To complement the results in Fig. 3e, we quantify how much structure-based priors improve model evidence per subject, context and MEQ subgroup. Fig. S3 shows, for each subject, context and MEQ subgroup, the log-Bayes factor comparing the hierarchical empirical Bayes model inverted with the evidence-weighted prior-variance transformation against the uninformed model (using data from the psilocybin session). In addition, we report the full high MEQ effective-connectivity profiles, obtained as the sum of the low-MEQ matrices (Fig. 3f–i) and the corresponding difference at high MEQ (Fig. 3j–m). Fig. S4 displays these unthresholded psilocybin profiles with self-connections converted back to the linear scale.

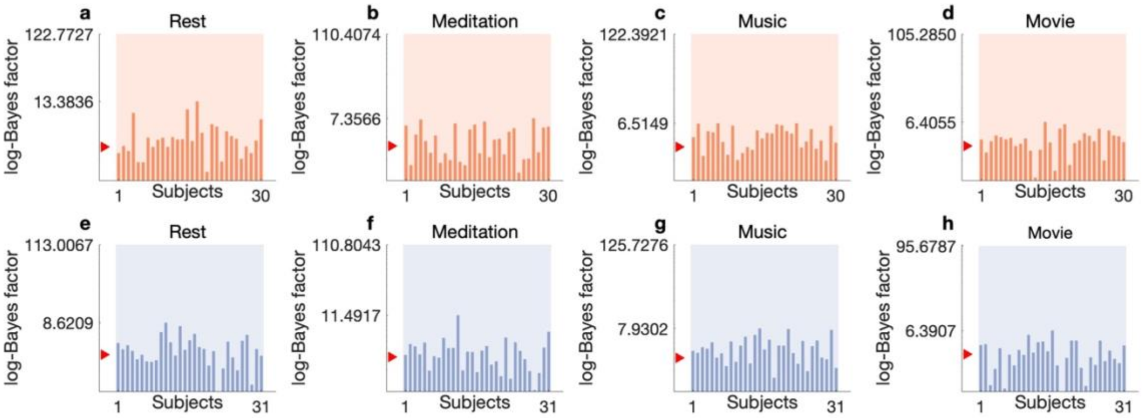

**Fig. S3 | Subject-level evidence gain from structure-based priors among mystical experience subgroups.** Panels show log-Bayes factors for each context: resting state (**a, e**), guided meditation (**b, f**), music listening (**c, g**), and movie viewing (**d, h**). Results shown for subjects that scored below (orange) (**a–d**) or above (blue) (**e–f**) the median on the revised Mystical Experience Questionnaire (MEQ). Opaque bars are per-subject log-Bayes factors comparing the hierarchical empirical Bayes model inverted with the evidence-weighted prior-variance transformation against the uninformed model. The semi-transparent background bar (spanning the panel width) is the group-level log-Bayes factors for the same comparison. The y-axis is on a log scale; tick labels mark the largest subject-level log-Bayes factors (lower tick) and the group-level log-Bayes factors (upper tick). The red triangle indicates log-Bayes factors of 3, a conventional benchmark for strong evidence. Higher values indicate stronger support for structure-based priors.

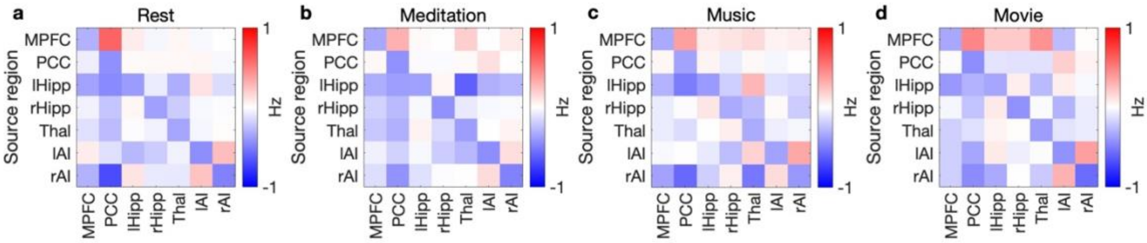

**Fig. S4 | Full psilocybin effective-connectivity profiles (unthresholded) for high mystical experience subgroup.** For each context—resting state (**a**), guided meditation (**b**), music listening (**c**), and movie viewing (**d**)—matrices depict the sum of psilocybin-modulated group-level effective connectivity for the low mystical experience subgroup (Fig. 3f–i) and the differences in effective connectivity for the high mystical experience subgroup (Fig. 3j–m): the full high mystical experience subgroup profile. Off-diagonal entries are directed inter-regional influences (Hz). Diagonal (self-connection) parameters have been converted from the log-normal parameterization back to the linear scale.

#### Mystical experience-independent psilocybin profiles and leave-one-out predictive checks

To complement Fig. 4, we provide two summaries. First, we show the psilocybin-session group mean of effective connectivity independent of MEQ (the intercept term when MEQ is included as a covariate, Fig. S5). Second, we provide leave-one-out posterior-predictive visualizations for MEQ prediction from selected left-hippocampal efferents. In all cases, models used the evidence-weighted, context-specific prior-variance transformations estimated earlier (Fig. 2); however, in the context of these analyses, structure-based priors exerted edgewise shrinkage (Methods).

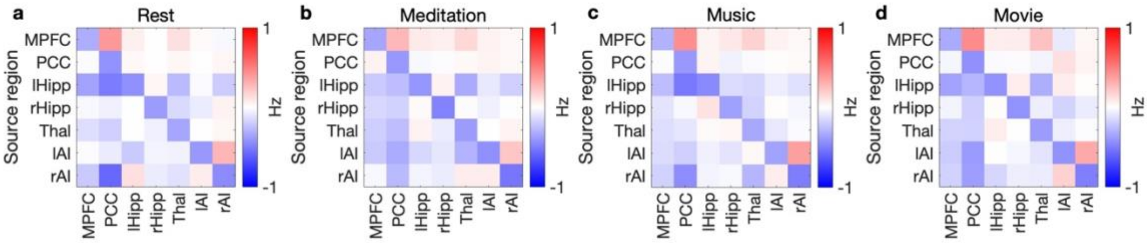

**Fig. S5 | Mystical experience-independent effective-connectivity profiles (unthresholded) under psilocybin.** For each context—resting state (**a**), guided meditation (**b**), music listening (**c**), and movie viewing (**d**)—matrices depict the psilocybin-session group mean of effective connectivity independent of mystical experience: the equivalent of the intercept term. Off-diagonal entries are directed inter-regional influences (Hz). Diagonal (self-connection) parameters have been converted from the log-normal parameterization back to the linear scale.

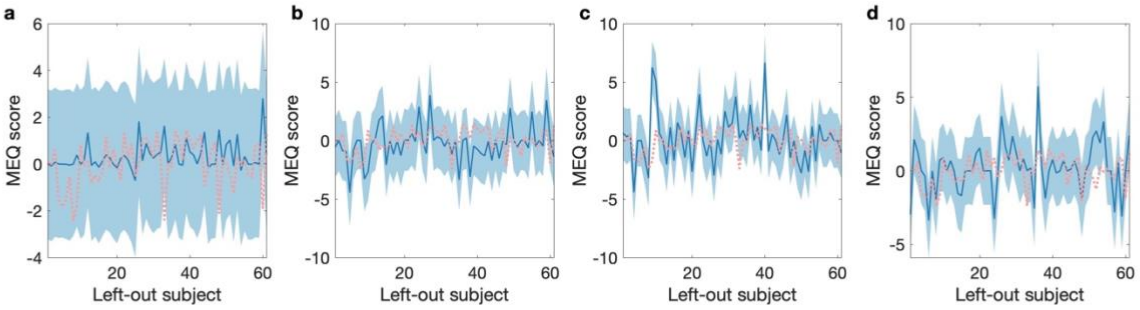

**Fig. S6 | Leave-one-out posterior-predictive checks for mystical experience from left-hippocampal efferents.** For each context—resting state (**a**), guided meditation (**b**), music listening (**c**), and movie viewing (**d**)—and the corresponding left-hippocampal efferent analyzed in Fig. 4f–i, the x-axis indexes the left-out subject. The blue line is the posterior mean prediction of that subject's z-scored Mystical Experience Questionnaire (MEQ) score, and the shaded band is the 90% credible interval (derived from the fold-specific predictive covariance). The orange dotted trace shows the observed z-scored MEQ. These panels visualize the same predictions summarized by in Fig. 4, making the uncertainty of each fold explicit.

#### Linking effective-connectivity variance to variability of neural signals and functional connectivity

Here, we illustrate how changes in the variance of effective connectivity translate to variability in observed BOLD signals and in model-free functional connectivity. With reference to Eq. 2, we simulated $S = 60$ synthetic subjects for a 7-region network ($n = 7,\ p = n^2$), across a linearly spaced vector of effective-connectivity standard deviations $\sigma_{\mathrm{E}} \in [0.10, 0.25]$ (20 samples). We constructed a third-level prior over group-level effective connectivity according to:

$$\boldsymbol{\theta}^{(2)} \sim \mathcal{N}\left(\boldsymbol{\mu}^{(3)}, \boldsymbol{\Sigma}^{(3)}\right), \qquad \boldsymbol{\Sigma}^{(3)} = \mathrm{diag}\left(\mathrm{vec}(\boldsymbol{V})\right), \tag{S11}$$

where $\boldsymbol{V} \in \mathbb{R}^{n \times n}$ assigns variance $\sigma_E^2$ to off-diagonal entries and a small, fixed variance to diagonals (self-connections):

$$v_{i,j} = \begin{cases} \sigma_E^2, & i \neq j \\ \sigma_{self}^2, & i = j \end{cases}, \text{ where } \sigma_{self}^2 = 1/64. \tag{S12}$$

Here, prior means $\boldsymbol{\mu}^{(3)}$, were consistent with those used in all analyses (Table S1). The second level followed Eq. 2 with an intercept-only design, and random effects parametrized according to a single precision component (Eq. S9).

For each $\sigma_E^2$, we sampled random variates from the model, leveraged subject-level effective connectivity matrices to generate neuronal time series ($T = 500$ timepoints) using a first-order multivariate autoregressive (MVAR) process (a discrete state equation, Eq. 1) with i.i.d. endogenous fluctuations of variance $1/64$, and convolved each regional trace with a canonical double-gamma HRF to obtain synthetic BOLD responses. From these, we computed both BOLD variability—per-subject regional standard deviations, averaged across regions and then across subjects—and functional connectivity variability: standard deviation across functional connectivity (Pearson-correlation) edges per subject, then mean across subjects.

Both BOLD and functional connectivity increased monotonically but nonlinearly with $\sigma_E^2$ (Fig. S7). Results are consistent with the fact that the state covariance of an MVAR process grows superlinearly as the effective connectivity matrix approaches instability (see discussion of Lyapunov relation in Friston and colleagues[30]), and for BOLD responses and functional connectivity, there are further nonlinearities from HRF convolution and correlation. Thus, we caution against interpreting variance changes in signals or functional connectivity as linearly proportional to variance changes in effective connectivity. MATLAB code that reproduces Fig. S7 is provided in the repository.

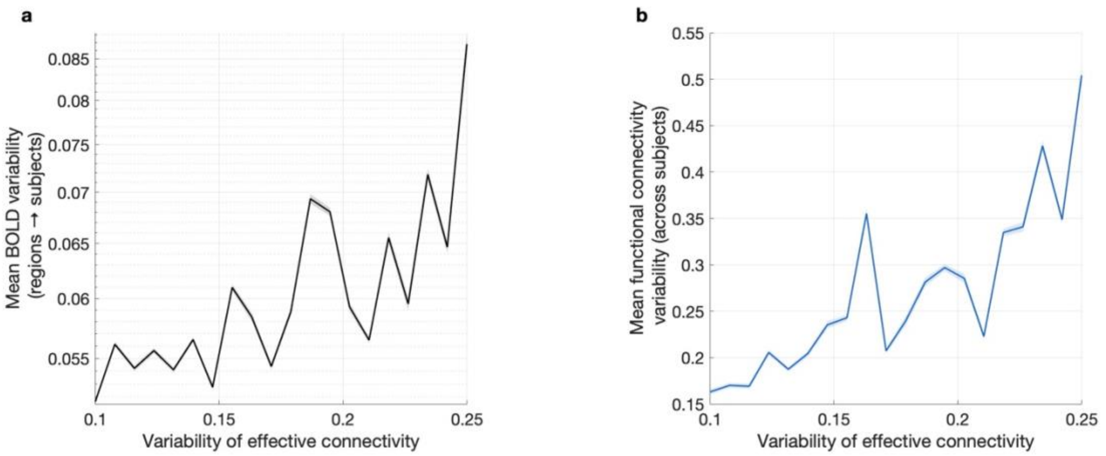

**Fig. S7 | Simulated linkage of effective-connectivity variance to variability of neural signals and functional connectivity.** (**a**) Mean (synthetic) blood-oxygen-level-dependent (BOLD) response variability (regional standard deviation averaged across regions then subjects) as a function of the standard deviation of off-diagonal effective-connectivity entries ($\sigma_{\mathrm{E}}$). (**b**) Mean functional connectivity variability (standard deviation across correlation edges per subject, then averaged across subjects) versus $\sigma_{\mathrm{E}}$. Each point reflects $S = 60$ synthetic subjects; lines show means with shaded areas showing its standard error.
